# Omicron: A heavily mutated SARS-CoV-2 variant exhibits stronger binding to ACE2 and potently escape approved COVID-19 therapeutic antibodies

**DOI:** 10.1101/2021.12.04.471200

**Authors:** Masaud Shah, Hyun Goo Woo

**Author notes:** Corresponding author, Hyun Goo Woo, M.D., Ph.D.

## Abstract

The new SARS-CoV-2 variant of concern “Omicron” was recently (Nov. 24th. 2021) spotted in South Africa and already spread around the world due to its enhanced transmissibility. The variant became conspicuous as it harbors more than thirty mutations in the spike protein with 15 mutations in the RBD region alone, potentially dampening the potency of therapeutic antibodies and enhancing the ACE2 binding. More worrying, Omicron infections have been reported in individuals who have received vaccines jabs in South Africa and Hong Kong. Here, we investigated the binding strength of Omicron with ACE2 and seven monoclonal antibodies that are either approved by FDA for COVID-19 therapy or undergoing phase III clinical trials. Computational mutagenesis and binding free energies could confirm that Omicron Spike binds ACE2 stronger than prototype SARS-CoV-2. Notably, three substitutions, i.e., T478K, Q493K, and Q498R, significantly contribute to the binding energies and doubled electrostatic potential of the RBD^Omic^-ACE2 complex. Instead of E484K substitution that helped neutralization escape of Beta, Gamma, and Mu variants, Omicron harbors E484A substitution. Together, T478K, Q493K, Q498R, and E484A substitutions contribute to a significant drop in the electrostatic potential energies between RBD^Omic^-mAbs, particularly in Etesevimab, Bamlanivimab, and CT-p59. CDR diversification could help regain the neutralization strength of these antibodies; however, we could not conduct this analysis to this end. Conclusively, our findings suggest that Omicron binds ACE2 with greater affinity, enhancing its infectivity and transmissibility. Mutations in the Spike are prudently devised by the virus that enhances the receptor binding and weakens the mAbs binding to escape the immune response.

## Introductions

Since the emergence of SARS-CoV-2 has been found to continuously evolve and raise new variants of concerns (VOCs) to avoid host hostilities, i.e., evade the host immune response, increase transmission, and aggress the pathogenesis of COVID-19. This host adaptation by the virus has been demonstrated by the rise of VOCs, including Alpha, Beta, Gamma, and Delta variants that weaken the neutralizing efficacy of antibodies (1–4). Most recently, a new strain of the SARS-CoV-2 named Omicron by the World Health Organization was recently (Nov. 24^th,^ 2021) emerged in South Africa and spread around the world within a short period.

Omicron harbors many novel mutations in both structural and non-structural proteins. The level of mutations that have led to serious concerns over vaccine failure, immune escape (5), and increased transmissibility have not been previously reported in any other VOC of the SARS0-CoV-2. More than 32 mutations were found in the Spike protein alone, where 15 of these mutations reside in the receptor-binding domain (RBD), which were vital to both receptor binding viral neutralizing antibodies. The non-structural proteins encoded by the ORF1ab contain mutations in the nsp3 (K38R, V1069I, Δ1265, L1266I, A1892T), nsp4 (T492I), nsp5 (P132H), nsp6 (Δ105-107, A189V), nsp12 (P323L), and nsp14 (I42V). Nsp3 (Plpro) and nsp5 (3Clpro, main protease) are proteases that cleave the polypeptide encoded by ORF1a and ORF1ab. 3Clpro and nsp12 (RNA-dependent RNA-polymerase (RdRp)) remained priority drug targets to block the polypeptide cleaving and viral protein synthesis (6). Using structural models, we found that mutations in the nsp5 and nsp12 are not close to the active site and may not hinder the effect of antiviral drugs; nonetheless, these proteins play a vital role in innate immune response (interferon induction) and requires further experimental investigation (6). In addition, Omicron harbors mutations in the other structural proteins, including Envelope (E) (T9I), Membrane (M) (D3G, Q19E, and A63T), and Nucleocapsid (N) (P13L, Δ31-33, R203K, G204R), which could further enhance the infectivity of the new variant. Since N protein is highly immunogenic (7, 8), these mutations could help escape the host immune response.

Omicron stood out as multiple mutations in the Spike protein are related to increased infectivity and antibody evasion. Out of 32 mutations, half hold the potential to dampen therapeutic antibodies’ potency and enhance the ACE2 binding. Besides, Omicron can infect vaccinated people as individuals who have already received mRNA-1273, AZD1222, and Ad26. In addition, COV2.S (VAC31518) jabs in South Africa and Hong Kong are infected.

We conducted molecular modeling and mutational analyses to understand how the new variant has enhanced its transmissibility and perhaps it may escape the FDA-approved Spike-neutralizing COVID-19 therapeutic antibodies. Seven therapeutic antibodies, including Etesevimab, Bamlanivimab, AZD8895, AZD1061, Imdevimab, Casirivimab, and CT-p59, were selected to investigate the RBD^Omic^ resistance to these antibodies. We observed that in addition to other mutations, T478K, Q493K, Q498R, and E484A substitutions mainly contribute to a significant drop in the electrostatic potential energies between RBD and antibodies. In addition, we found that Omicron Spike binds ACE2 stronger than prototype SARS-CoV-2; particularly, three substitutions, i.e., T478K, Q493K, and Q498R, significantly contribute to the binding energies and doubles electrostatic potential of the RBD-ACE2 complex.

## Results

### Mutations in the Omicron RBD strengthen the Spike-ACE2 interaction

Mutations in the Spike and other genes make Omicron unique among previously reported SARS-CoV-2 VOCs. According to the unrooted phylogeny constructed from the global ~4000 full-genome SARS-CoV-2 sequences submitted at Global initiative on sharing all influenza data (GISAID), Omicron stands distant from other VOCs (**Figure 1A**). A full-length trimeric 3D model was constructed by substituting the respective amino acids of previously reported reference (Wuhan strain, PDB ID: 7VNE) structure into Omicron. There are three deletion sites in the N-terminus domains (NTD) and at least 15 substitutions in the RBD region. Omicron Spike also harbors mutations such as K417, T478, E484, and N501 that are reported in previous VOCs **(Figure1B**). At least eleven out of fifteen mutated residues are involved in ACE2 binding and substantially affect its binding affinity (**Figure 1C**). In addition, compared to the prototype SARS-CoV-2, Omicron Spike has three deletions, i.e., Δ69-70, Δ143-145, and Δ211, and one highly charged insertion at 214 positions in the Spike, i.e., ins214EPE.

**Figure 1:**
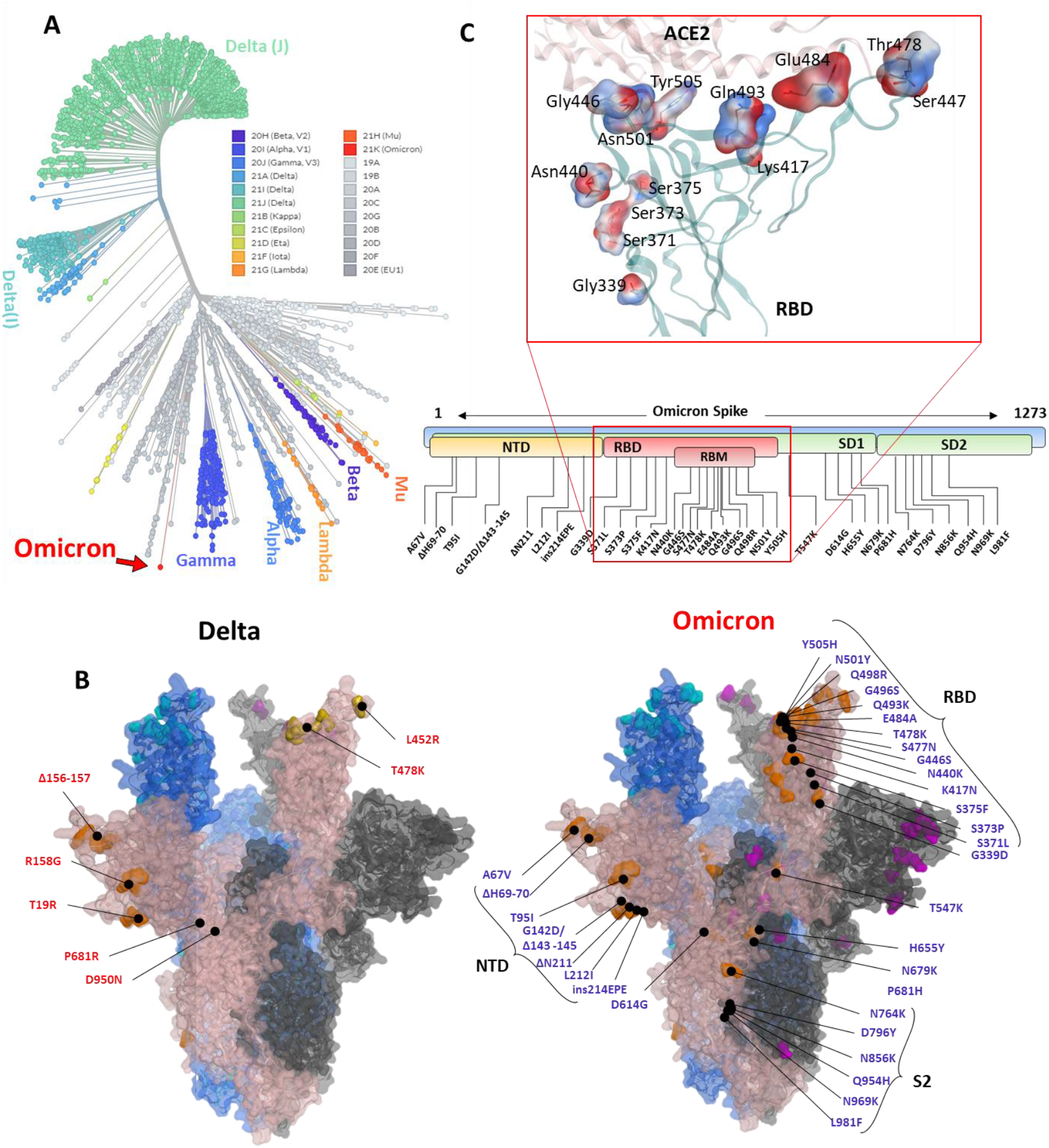
Phylogeny of the Omicron and annotation of the mutation in Spike protein. The Unrooted phylogenetic tree was constructed from the Nextstrain servers. Wuhan-Hu-1/2019 strains were taken as a reference sequence. B) The full-length Delta and Omicron Spike were built to annotate the relative (not exact) positions of the mutations on the surface map of Spike. C) The amino acids mutated in the RBD of Omicron are shown concerning the ACE2 interface. Residues are colored according to the electrostatic map of the WT strain. Respective Omicron mutations are depicted in the panel below the RBD surface map.

To monitor the relative binding strength of RBD-ACE2 complexes of both prototype and Omicron strains, we used a protein design strategy and calculated binding affinity and stability changes. Although individually substituted residues had a slight effect on the local stability of the RBD-ACE2 complexes (**Figure 2A**), a substantial increase in the binding affinity by T478K, Q493K, and Q498R overridden this effect and led to an overall increase (ΔG^WT^=64.65 kcal/mol<ΔG^Omic^=83.79 kcal/mol) in the binding affinity of the RBD^Omic^ with ACE2, as investigated through endpoint MM/GBSA (**Figure 2B**). Since five residues in the RBM region of RBD are mutated from polar to positively charged residues, i.e., N440K, T478K, Q493K, Q498R, and Y505H, we investigated the change in electrostatic potential of the RBD^Omic^ relative to that of RBD^WT^. Surprisingly, we could see that the electrostatic energy of ACE2-RBD^Omic^ was double as that of ACE2-RBD^WT^, which in turn doubled the Polar Solvation free energies of the ACE2-RBD^Omic^ (**Figure 2B**). Per residues, energy distribution suggests that mutations in the RBD^Omic^ directly participate in binding and enhance the binding strength of amino acids in the same network (**Figure 2C**). Among 15 substituted amino acids, K417N and Y505H exhibited a slight reduction in binding energy due to the breakage of salt bridges between K417 of the RBD and D30 of ACE2; nonetheless, this breakage was compensated by the salt bridge between E35 of ACE2 and Q493K substitution in RBD^Omic^ (**Figure 2C,** right panel). Together, this data suggests that Omicron binds ACE2 with greater affinity, partly explaining its increased transmissibility. Although RBD^Omic^ exhibits a higher binding affinity to ACE2, it is hard to predict the pathogenicity of the new strain for this data.

**Figure 2:**
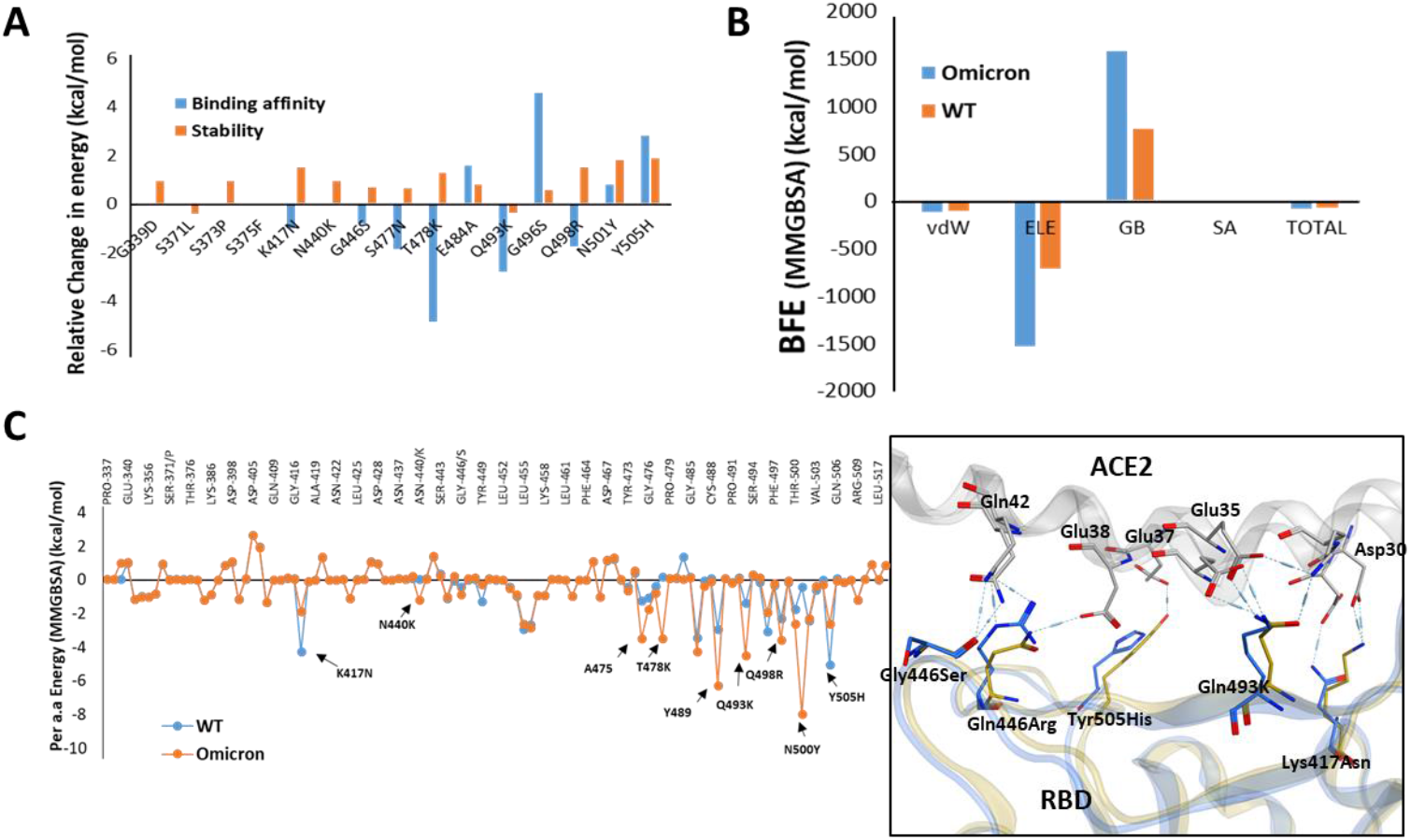
Relative effect of mutations in Omicron RBD on the ACE2 binding. **A**) Effect of 15 individual mutations on the binding and stability of RBD^Omic^-ACE2 was monitored relative to that of RBD^WT^-ACE2. **B)** The overall binding energies (measured through endpoint MM/GBSA) as consequences of all 15 mutations at once were monitored for both RBD^Omic^-ACE2 and RBD^WT^-ACE2. **C)** Per-residue change in the binding affinity was monitored, and the hotspots of RBD were labeled. The change in the hydrogen bonds network of the elected hotspots is shown at the right.

### Mutations in the RBD^Omic^ deteriorate the binding of therapeutic antibodies and garble their epitopes on RBD

To find how these mutations in the RBD that strengthen the RBD^Omic^-ACE2 interaction will affect the RBD-targeting COVID-19 therapeutic antibodies, we constructed structural models of seven mAbs bound to RBD^Omic^ (**Figure 3A-D**). Only one mAb (CT-p59) formulated by Celltrion is used as solo COVID-19 therapeutic and undergoing phase III clinical trials (9), whereas the rest of six mAbs are approved for COVID-19 therapies on an emergency basis (10). To tackle the immune escape by rising mutants, these antibodies are used as cocktail therapy (mAbs cocktails are devised by the companies which sponsor the drugs as indicated in **Figure 3A-D**).

**Figure 3:**
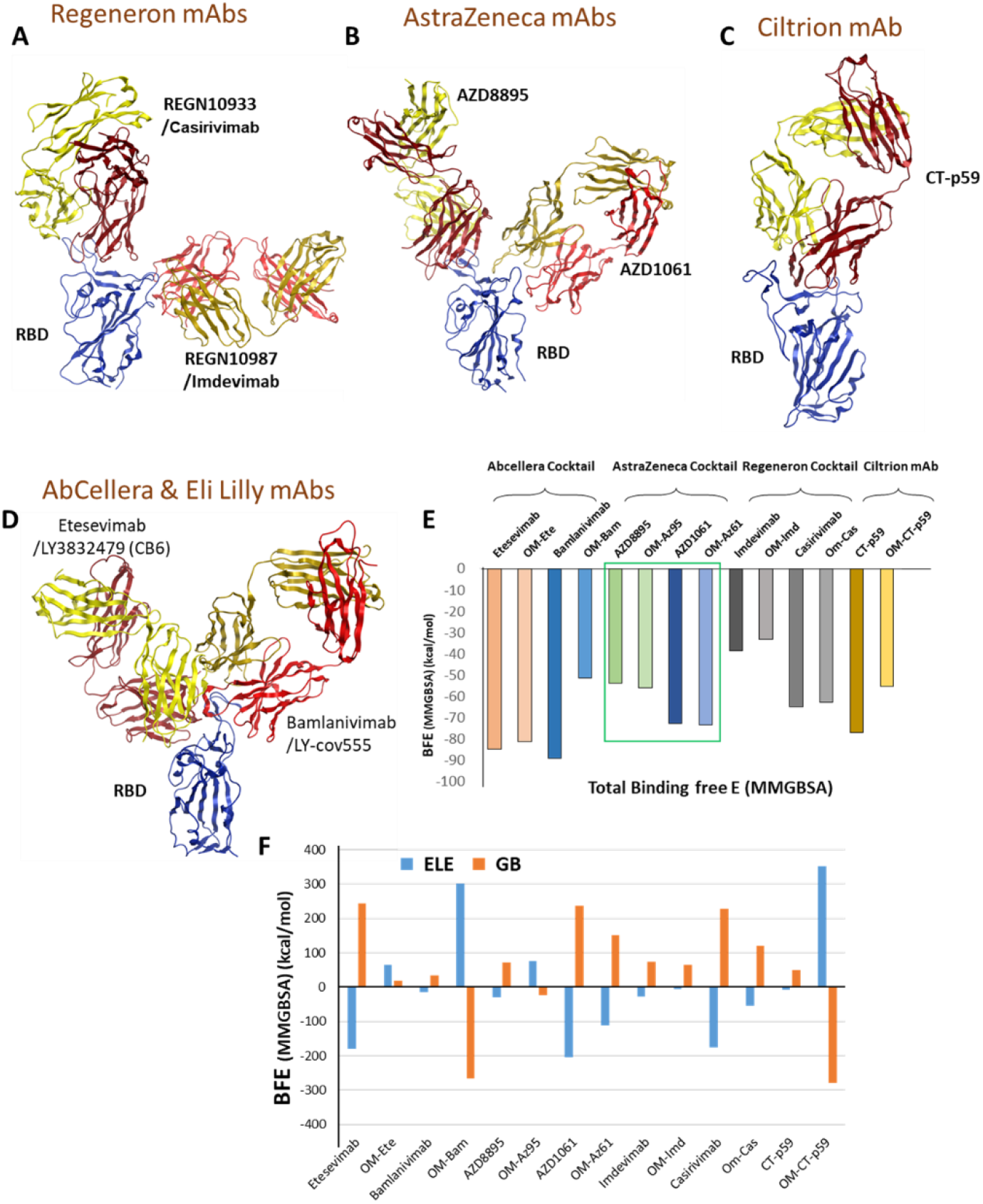
Mutations in the Omicron RBD distort the epitopes of therapeutic mAbs. **A-D)** Crude epitopes of seven selected mAbs are shown on the RBD. Antibodies used as cocktails are labeled with their sponsors. All variable light chains are colored yellow or orange and variable heavy chains are colored red. **E)**Changes in the binding affinity of the RBD^Omic^-mAbs relative to RBD^WT^-mAbs are shown. The Binding energies were calculated through endpoint MM/GBSA). **F)** Changes in the electrostatic potentials and polar solvation energies are shown to each RBD-mAb complex.

Since these mAbs are not sharing overlapping epitopes on the RBD, except Etesevimab and Bamlanivimab (sponsored by AbCellera), where the light chain variable domains make a slight clash, we investigated the change in their interface and binding strength with RBD^Omic^ individually. Surprisingly, we found a substantial drop in the total binding energies of Bamlanivimab and CT-p59 when bound to RBD^Omic^ (**Figure 3E**). The total binding energy is the sum of four energies (listed in Table 1). We could see that vdW (Van der Waals potentials) and SA (solvation free energy) energies did not affect the binding strength; however, electrostatic potentials (ELE) had a significant shift in the calculated energies. The magnitude of the change in ELE energies was similar to that we observed in RBD^Omic^-ACE2 case, except that the effect was opposite, i.e., in RBD^Omic^-ACE2 ELE energies favor the binding whereas RBD^Omic^-mAbs ELE opposed the binding strength (**Figure 3F**). Except AZD1061 (AstraZeneca) where the ELE energies slightly dropped (RBD-AZD1061 = −204.24 kcal/mol > RBD^Omic^-AZD1061 = −112.35 kcal/mol) other mAbs showed a significant drop.

**Table 1:** The binding energies of RBD^Omic^ and RBD^WT^ with seven therapeutic mAbs are listed.

| Sponsor | mAbs | VDW | ELE | GB | SA | TOTAL |
| --- | --- | --- | --- | --- | --- | --- |
| **AbCellera & Eli Lilly** | Etesevimab | -131.41 | -179.48 | 243.28 | -17.15 | -84.76 |
|  | Omic-Ete | -147.42 | 65.04 | 19.31 | -18.16 | -81.24 |
|  | Bamlanivimab | -95.81 | -13.97 | 33.7 | -13.01 | -89.1 |
|  | Omic-Bam | -108.74 | 602.59 | -532.27 | -13.12 | -51.54 |
| **AstraZeneca** | AZD8895 | -84.37 | -30.35 | 71.27 | -10.46 | -53.92 |
|  | Omic-As95 | -96.77 | 76.73 | -24.25 | -11.62 | -55.91 |
|  | AZD1061 | -94.96 | -204.24 | 238.04 | -11.49 | -72.64 |
|  | Omic-As61 | -98.83 | -112.35 | 149.92 | -12.29 | -73.55 |
| **Regeneron** | Imdevimab | -76.1 | -27.36 | 74.73 | -9.67 | -38.4 |
|  | Omic-Imd | -81.32 | -5.84 | 64.23 | -10.15 | -33.08 |
|  | Casirivimab | -103.52 | -174.81 | 228.04 | -14.66 | -64.94 |
|  | Omic-Cas | -113.25 | -55.14 | 119.9 | -14.42 | -62.91 |
| **Celltrion** | CT-p59 | -105.66 | -7.38 | 49.61 | -13.75 | -77.18 |
|  | Omic-CT-p59 | -114.85 | 352.19 | -278.48 | -14.27 | -55.4 |

mAbs monoclonal antibodies: VDW, Van der Waals potentials: ELE, electrostatic potentials: GB, polar solvation potentials (generalized born model): SA, Nonpolar contribution to the solvation free energy calculated by an empirical model.

To understand which mutations are particularly involved in weakening the RBD^Omic^-mAbs interactions, per residues, change in energies were calculated for CT-p59 and Bamlanivimab when bound RBD^WT^ and RBD^Omic^. Two hotspots, R96 in CDRL3 and R50 in CDRH2 of the Bamlanivimab, which establish highly stable salt bridges with the E484 of RBD^WT^, completely lost their binding upon E484A mutation in RBD^Omic^ (**Figure 4A**). In addition, E102 and R104 in CDRH3 showed a 50% reduction in binding energies. Similarly, the hotspots in CDRL1 and CDRH3 of CT-p59 entirely or partly lost their bindings due to E484A, Q493K, and Y505H mutations in RBD^Omic^. One mutation, i.e., N501Y, slightly favors the binding energy upon establishing a hydrogen bond with D57 in CDRH2 (**Figure 4B**). Overall, these data suggest that mutations in the Omicron spike were precisely engineered to utilize the same mutations to enhance receptor binding and resist antibody binding. This raises very serious concerns about the efficacy of therapeutic mAbs in Omicron patients.

**Figure 4:**
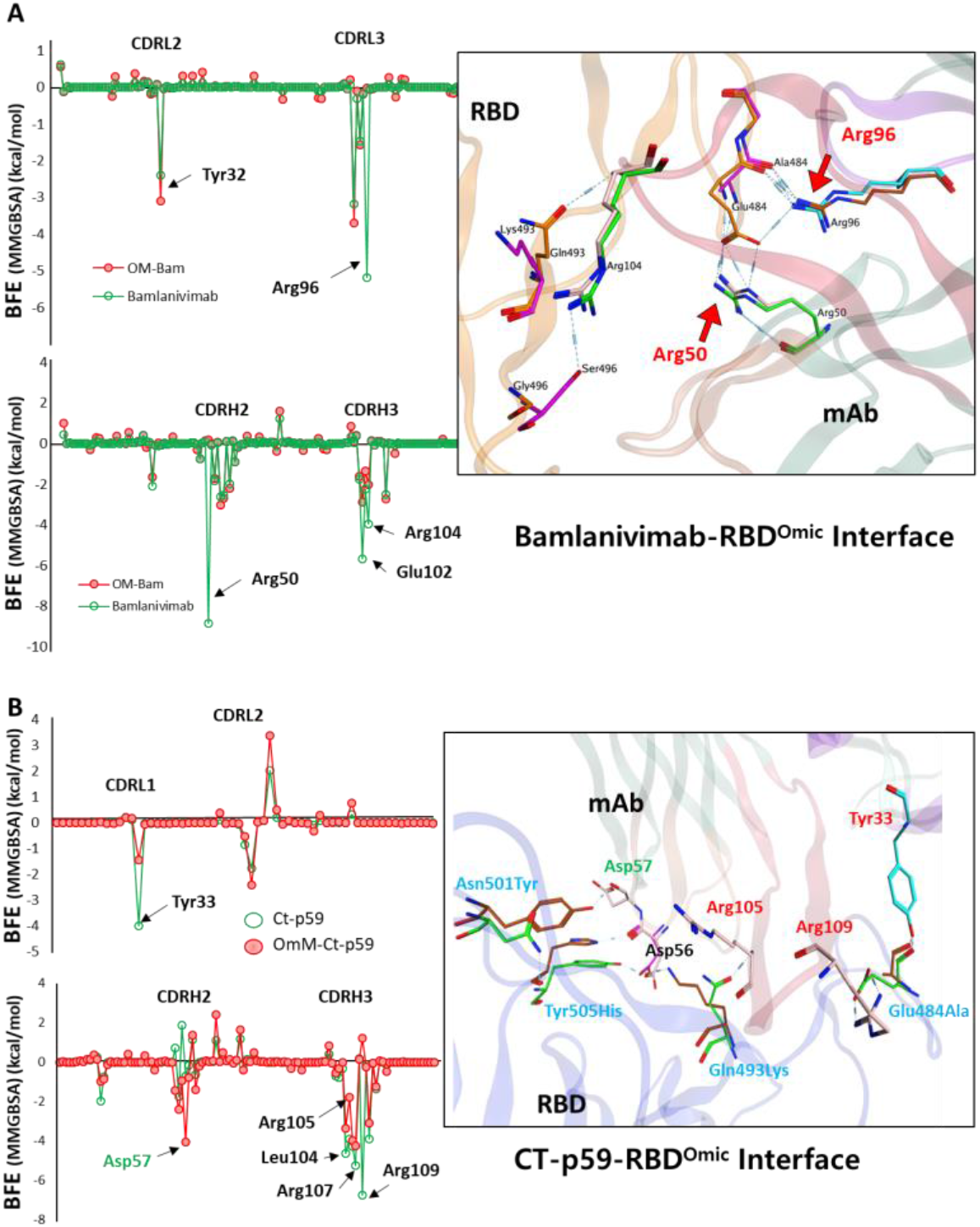
Per-residue changes in the binding affinity of RBD-mAbs were monitored, and the hotspots on CDRs of **(A)** Bamlanivimab and **(B)** CT-p59 are labeled. The change in the hydrogen bonds network of the selected hotspots is shown at the right.

## Discussion

To this end, this fact is well established that SARS-CoV-2 is rapidly evolving and makes at least two mutations per month in its genome globally (11, 12). The virus’s capability to adopt the host environment by increasing transmissibility and evading immune response, as exemplified by the continuous rise of variants of concern, including alpha, Beta, Gamma, and Delta (13, 14). Although tremendous efforts have been made in vaccine development and COVID-19 therapeutics, including mAbs and COVID-19 pills by Merck, the continuous emergence of VOCs has raised concerns over the efficacy of neutralizing antibodies induced by the vaccines, collected from the convalescent plasma or developed otherwise (14, 15). Even though these variants have a limited number of mutations, although delirious, they successfully escape the immune response, at least partly is not entirely. Omicron harbors four to five times more mutations in its spike protein than other SARS-CoV-2 VOCs (**Figure 1B**). Nearly half of these mutations are within the RBD region that is most antigenic and remained the main focus of Spike-targeting therapeutics antibodies. Researchers around the globe are racing to determine whether Omicron poses a threat to the immunity induced by the COVID-19 vaccine (5).

Till now, there is no data available that could show the response of this heavily mutated Spike towards therapeutic mAbs and change in its binding affinity with ACE2. Thus, we used the previously available structural data of Spike RBD-binding antibodies, Spike itself, and Spike-ACE2 complexes and constructed the mutated Spike of Omicron. Omicron Spike contains some of the mutations reported in previous VOCs, particularly D614, that enhances the receptor-binding by increasing its “up” conformation and the overall density of Spike protein at the virus’s surface (16, 17). In addition, five of the amino acids within the RBD region are mutated from polar to positively charged residues (K, R, or H) that paradoxically enhance the receptor binding and weaken the Spike neutralizing interactions (**Figure 2C and Figure 4**). Among all seven antibodies that are investigated here, we speculate that AZD1061 and Casirivimab may moderately hold their Omicron neutralization, whereas the rest may not (**Figure 3F**).

The variant was named from B.1.1.529 to Omicron by WHO on Nov. 24th, 2021, whereas the strain was spotted by the researchers from genome-sequencing of the SARS-CoV-2 in Botswana and researchers from researchers the University of KwaZulu-Natal, South Africa, between 12 and 20 November. This indicates that the strains had already appeared at the first week of November and spreads into another part of the world through travelers throughout the whole month. Hence, Omicron’s threat is global now, and it is also quite clear that the new variant is more transmissible than Delta. Therefore, it is of utmost priority to follow the variant more closely and speed up the vaccination process as it significantly drops the COVID-19 infection VOCs including Delta (18).

## Methods

### Materials used in this project

For full-length trimeric Spike a previously reported PDB ID: 7VNE was used to rebuild Omicron spike protein using a Swiss-Model server (19). Other structures used in this study are listed below: RBD-ACE2 (PDB ID: 6MOJ), RBD-Etesevimab (PDB ID: 7C01), RBD-Bamlanivimab (PDB ID: 7KMG), RBD-CT-p59 (PDB ID: 7CM4), RBD-AZD1061 (PDB ID: 7I7E), RBD-AZD8895 (PDB ID: 7I7E), RBD-Casirivimab (PDB ID: 6XDG), RBD-Imdevimab (PDB ID: 6XDG).

### Computational tools used in this project

For protein structure visualization VMD (20), Pymol (https://pymol.org), and Chimaera Chimera (21) packages were used. For electrostatic surfaces, isolation of the proteins, APBS and APBSrun plugins in Pymol and VMD were utilized. The interfaces of RBD^WT^ and RBD^Omic^ with mAbs and ACE2 were analyzed by online server PDBePISA (v1.52) (22), and the binding contribution of individual amino acids was determined. For constructing mutant RBD, free BIOVIA Discovery Studio Visualizer was used (http://www.accelrys.com). All complexes were solvated with TIP3P water cubic box of dimension boundaries extended to 10 Å from protein atoms and neutralized with counter ions, Na+/Cl-, wherever needed. The neutralized systems were energy minimized in GROMACS 2019.6 (23) using CHARMM37 force field (24) and steep descent algorithm. For endpoint binding free energies calculations, the HawkDock server was utilized (25). The hotspot results were validated through the DrugScorePPI web server (26). The unrooted phylogenetic tree was constructed from the Nextstrain (27) servers using ~4000 full-length SARS-CoV-2 sequences from GISAID (28) database with reference to Wuhan-Hu-1/2019 as a reference sequence.

## Authors’ contributions

M.S. and H.W. contributed toward conceptualization of the project and designed the methodology. M.S. and H.W. wrote the original manuscript draft; H.W supervised the study and provided funding acquisition.

